# Reference genome for the Northern bat (*Eptesicus nilssonii*), a most northern bat species

**DOI:** 10.1101/2023.07.18.549444

**Authors:** Veronika N. Laine, Arto T. Pulliainen, Thomas M. Lilley

## Abstract

The northern bat (*Eptesicus nilssonii*) is the most northern bat species in the world. Its distribution covers whole Eurasia, and the species is thus well adapted to different habitat types. However, recent population declines have been reported and rapid conservation efforts are needed. Here we present a high-quality *de novo* genome assembly of a female northern bat from Finland (*BLF_Eptnil_asm_v1*.*0*). The assembly was generated using a combination of Pacbio and Omni-C technologies. The primary assembly comprises 726 scaffolds spanning 2.0 Gb, represented by a scaffold N50 of 102 Mb, a contig N50 of 66.2 Mb, and a BUSCO completeness score of 93.73%. Annotation of the assembly identified 20,250 genes. This genome will be an important resource for the conservation and evolutionary genomic studies especially in understanding how rapid environmental changes affect northern species.

## Introduction

The northern bat belongs to the global and speciose genus of serotine bats, *Eptesicus* (family Vespertilionidae, subfamily Vespertilioninae). It exhibits a wide trans-continental distribution across Eurasia, running pretty much continuous across Siberia from Hokkaido Island to Fennoscandia (Suominen et al., 2020) (Figure 1). Because of its broad distribution range and several isolated relict populations, which have arisen as a consequence of distribution range shifts caused by the last ice age, the possibility of differentiation across populations is apparent, even to the degree of some isolated populations forming distinct subspecies. With a distribution that extends well over the Arctic circle (Kotila et al., 2022; Siivonen & Wermundsen, 2008) to the north, the species appears to be well adapted to northerly latitudes short active seasons (Kotila et al., 2022), short nights (Vasko et al., 2020) and a long season of inactivity during the winter (Blomberg et al., 2021).

**Figure 1.**
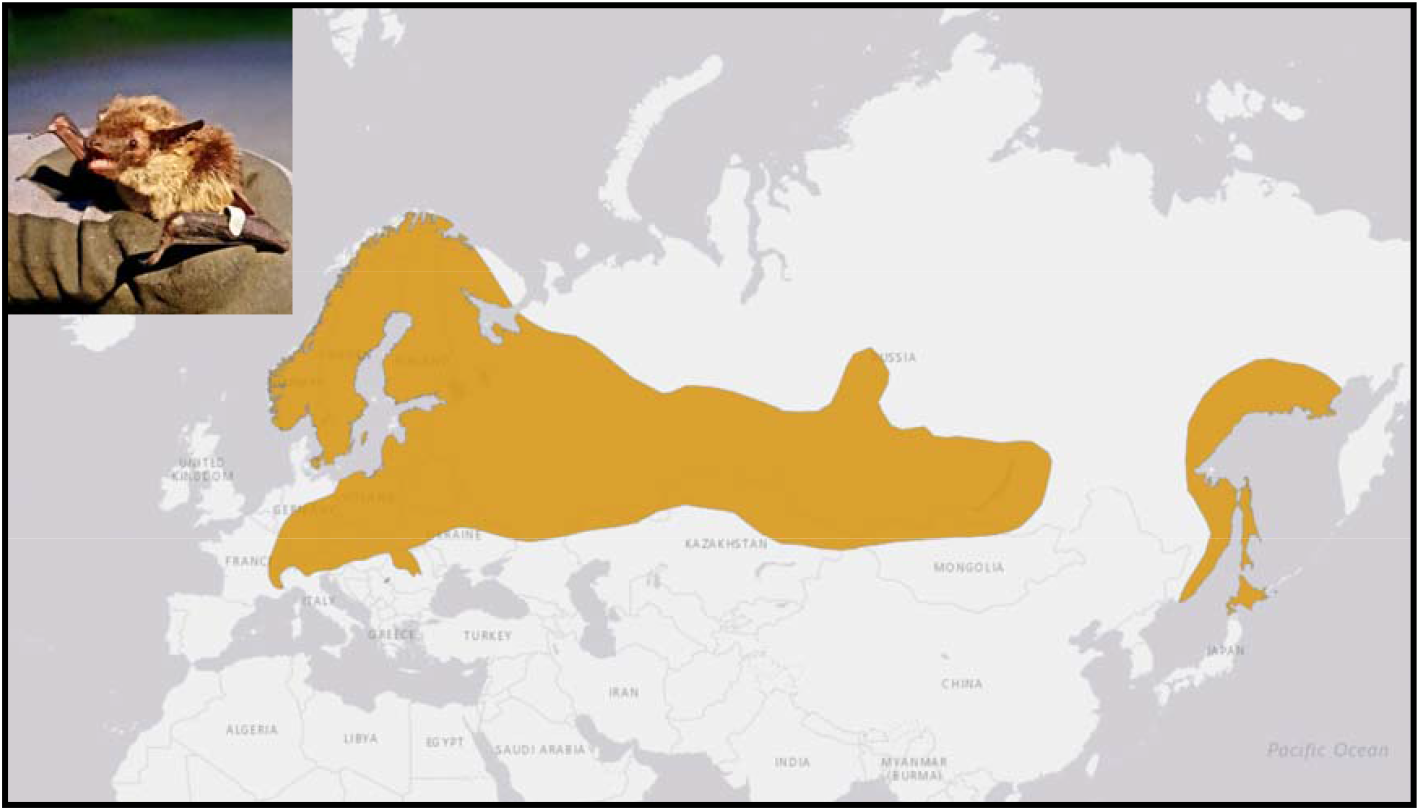
Distribution range of E. nilssonii. Map adapted from www.iucn.org.

However, due to the extreme latitudinal distribution of the species, it is particularly vulnerable to the effects of climate change. Estimates suggest that the Arctic and Sub-Arctic are experiencing the most rapid effects of climate change, which along with urbanization and modification of natural habitats puts wildlife under immeasurable pressure (Rantanen et al. 2022). Furthermore, the northern bat population in southern Sweden has already seen decline of c. 50%, which has been partially attributed to competition from bat species increasing their distribution range to the north as a consequence of climate change (Rydell et al., 2020). The production of a high-quality reference genome allows the investigation of the effects of environmental change on the northern bat to assist in better population viability assessments and planning of conservation measures via population genetic approaches.

## Methods

### Biological materials

A female northern bat was sampled in February 2013 from Lieto, Finland (60.57 N, 22.43 E). An active bat was found inside a house in the middle of the hibernation season. The unusually behaving bat subsequently died after it had been rescued. From this sample a cell clone was derived from primary cells isolated from bat kidney tissue and immortalized via the transfer and stable production of the Simian virus 40 Large T antigen (SV40LT). Kidneys were cut in small pieces (ca. 8 mm^3^) and incubated in 0.05% Trypsin-EDTA solution (Gibco # 25300-054) for 15 h at 4 ºC, followed by 1 h at 37 ºC in gentle rotation. Tissue pieces and detached cells were pelleted by centrifugation (300 x g, 10 min, room temperature), washed once with calcium- and magnesium free phosphate buffered saline (PBS, Lonza # BE17-516F), and resuspended in Dulbecco’s modified Eagle medium (DMEM, Lonza # 12-709F) supplemented with 10% heat-inactivated fetal bovine serum (iFBS, Gibco # 10270-106), 100 U /mL penicillin and 100 μg / mL streptomycin antibacterial mixture (PEN / STREP, Biochrom # A2213) and 2.5 μg /mL of anti-fungal amphotericin B (Sigma # 2942). The mixtures of tissue pieces and cells were incubated at 37 °C in humidified atmosphere with 5% CO_2_ in 6-well cell culture plate format for up to 3 weeks. The medium was exchanged in every 3-4 days. The cells were detached by trypsinization, recultured in DMEM/ iFBS /PEN / STREP, and transfected (Fugene 6, Promega # E269A) with pBABE-puro SV40 LT plasmid (Addgene # 13970, selection with 4 μg /mL puromycin A111380-03), allowing ectopic expression of the SV40LT. The SV40LT-immortalized cells were serially diluted and cultured in 96-well cell culture plate format in DMEM /iFBS /PEN / STREP, allowing isolation of clonal cell lines originating from single transfected cells (Supplementary Figure 1). The clonal cell lines were expanded and routinely cultured in DMEM /iFBS /PEN /STREP at 37 °C in humidified atmosphere with 5% CO _2_. To prepare the sample for sequencing, 25 000 000 cells of the clonal isolate 8+ (Supplementary Figure 1) were collected by trypsinization, washed twice with PBS and then frozen for storage at –80°C. The high molecular weight DNA was extracted from the cell culture with Qiagen DNeasy Blood & Tissue Kit. DNA samples were quantified using Qubit 2.0 Fluorometer (Life Technologies, Carlsbad, CA, USA).

### DNA Sequencing and Genome Assembly

#### Pacbio library preparation and sequencing

The PacBio SMRTbell library (∼20kb) for PacBio Sequel was constructed using SMRTbell Express Template Prep Kit 2.0 (PacBio, Menlo Park, CA, USA) using the manufacturer recommended protocol. The library was bound to polymerase using the Sequel II Binding Kit 2.0 (PacBio) and loaded onto PacBio Sequel II. Sequencing was performed on PacBio Sequel II 8M SMRT cells generating 281.2 gigabases of data.

#### Dovetail Omni-C library preparation and sequencing

For each Dovetail Omni-C library, chromatin was fixed in place with formaldehyde in the nucleus and then extracted. Fixed chromatin was digested with DNAse I, chromatin ends were repaired and ligated to a biotinylated bridge adapter followed by proximity ligation of adapter containing ends. After proximity ligation, crosslinks were reversed, and the DNA purified. Purified DNA was treated to remove biotin that was not internal to ligated fragments. Sequencing libraries were generated using NEBNext Ultra enzymes and Illumina-compatible adapters. Biotin-containing fragments were isolated using streptavidin beads before PCR enrichment of each library. The library was sequenced as 150bp paired-end on an Illumina HiSeqX platform to produce an approximately 30x sequence coverage.

#### Assembly and scaffolding

For the Pacbio assembly, Wtdbg2 (version 2.5) (Ruan and Li, 2020) was run with the following parameters: --genome_size 2.0g --read_type sq --min_read_len 20000 --min_aln_len 8192 using the Pacbio CLR reads. Blobtools (version 1.1.1) (Laetsch and Blaxter) was used to identify potential contamination in the assembly based on BLAST (version 2.9) (Altschul et al. 1990) results of the assembly against the nt database. A fraction of the scaffolds was identified as contaminant and were removed from the assembly. The filtered assembly (filtered.asm.cns.fa) was then used as an input to purge_dups (version 1.1.2) (Guan et al. 2020) and potential haplotypic duplications were removed from the assembly, resulting in the final purged.fa assembly.

The input *de novo* assembly and Dovetail OmniC library reads were used as input data for HiRise, a software pipeline designed specifically for using proximity ligation data to scaffold genome assemblies (Putnam et al., 2016). Dovetail OmniC library sequences were aligned to the draft input assembly using bwa (0.7.17) (Li 2013) (https://github.com/lh3/bwa) only using reads with MQ>50. The separations of Dovetail OmniC read pairs mapped within draft scaffolds were analyzed by HiRise to produce a likelihood model for genomic distance between read pairs, and the model was used to identify and break putative misjoins, to score prospective joins, and make joins above a threshold. Due to possibility of remnants of plasmid containing SV40LT remaining in the reference genome, the sequence of SV40LT (NC_001669.1) was searched with Blast from the final assembly.

#### RNA sequencing

RNA was extracted from the cultured cells with QIAGEN RNeasy Plus Kit and a standard RNA library was prepared with rRNA-depletion with QIAGEN FastSelect HMR kit. The libraries were sequenced at an Illumina NovaSeq platform (Illumina, CA) targeting approximately 20 million 150 bp paired end reads.

#### Annotation

Repeat families found in the genome assemblies of *Eptesicus nilssonii* were identified de novo and classified using the software package RepeatModeler (version 2.0.1) (Flynn et al. 2020). RepeatModeler depends on the programs RECON (version 1.08) (Bao & Eddy 2002) and RepeatScout (version 1.0.6) (Price et al. 2005) for the de novo identification of repeats within the genome. The custom repeat library obtained from RepeatModeler were used to discover, identify and mask the repeats in the assembly file using RepeatMasker (Version 4.1.0) (Smit, Hubley & Green RepeatMasker at http://repeatmasker.org). Coding sequences from *Eptesicus fuscus* (GCF_000308155.1), *Myotis myotis* (GCF_014108235.1) and *Pipistrellus kuhlii* (GCF_014108245.1) were used to train the initial ab initio model for *Eptesicus nilssonii* using the AUGUSTUS software (version 2.5.5) (Stanke et al. 2008). Six rounds of prediction optimisation were done with the software package provided by AUGUSTUS. The same coding sequences were also used to train a separate ab initio model for *Eptesicus nilssonii* using SNAP (version 2006-07-28) (Korf 2004). RNAseq reads were mapped onto the genome using the STAR aligner software (version 2.7) (Dobin et al. 2013) and intron hints generated with the bam2hints tools within the AUGUSTUS software. MAKER (v3.01.03) (Cantarel et al. 2008), SNAP and AUGUSTUS (with intron-exon boundary hints provided from RNAseq) were then used to predict for genes in the repeat-masked reference genome. To help guide the prediction process, Swiss-Prot peptide sequences from the UniProt database were downloaded and used in conjunction with the protein sequences from *Eptesicus fuscus, Myotis myotis* and *Pipistrellus kuhlii* to generate peptide evidence in the Maker pipeline. Only genes that were predicted by both SNAP and AUGUSTUS softwares were retained in the final gene sets. To help assess the quality of the gene prediction, Annotation Edit Distance (AED) scores were generated for each of the predicted genes as part of the MAKER pipeline. Genes were further characterised for their putative function by performing a BLAST search of the peptide sequences against the UniProt database. tRNA were predicted using the software tRNAscan-SE (version 2.05) (Chan et al. 2021).

#### Mitochondrial genome assembly

The raw Pacbio subreads were aligned *Myotis myotis* mitochondria (NC_029346.1) with minimap2 (version 2.24) (Li 2018). The aligned reads were extracted with Samtools sort and transformed from bam-format to fastq-format with Samtools fastq (version 1.16.1) (Danecek et al. (2021). The mitochondrial genome was assembled with Canu (version 2.1.1.) (Koren et al. 2017) (genomeSize=20k, corOutCoverage=999) and visual inspection and consensus sequence of the assembled contigs was made with Geneious (version 11.0.3.) (https://www.geneious.com). Annotation was done with MitoZ (version 3.4) (Meng et al., 2019) using clade Chordata.

#### Genome quality assessment

BUSCO (version 4.0.5) (Manni et al. 2021) was used to evaluate genome quality and completeness with the eukaryotes database (eukaryota_odb10) that contains 255 genes and 70 taxa. See Table 1 for a list of software used in this study.

**Table 1.** Assembly pipeline and software usage.

| Method | Software | Version |
| --- | --- | --- |
| K-mer counting | Meryl | 1.4 |
| De novo assembly (contigs) | Wtdbg2 | 2.5 |
| Contamination screening |  |  |
| Alignment | Blast | 2.9 |
| Screening | Blobtools | 1.1.1 |
| Low-coverage, duplicated contigs removal | purge_dups | 1.1.2 |
| **Scaffolding** |  |  |
| Omni-C scaffolding | HiRise | n/a |
| Omni-C alignment to draft | bwa | 0.7.17 |
| Omni-C Contact map generation | Juicer | 1.5 |
| **Annotation** |  |  |
| Repeat family identification | RepeatModeler | 2.0.1 |
|  | RECON | 1.0.8 |
|  | RepeatScout | 1.0.6 |
| Repeat masking | RepeatMasker | 4.1.0 |
| Gene prediction optimisation | AUGUSTUS | 2.5.5 |
|  | SNAP | 2006-07-28 |
| RNA read alignment | Star | 2.7 |
| Intron hints | bam2hints (AUGUSTUS) | 2.5.5 |
| Annotation | MAKER | v3.01.03 |
|  | AUGUSTUS | 2.5.5 |
|  | SNAP | 2006-07-28 |
| Gene function prediction | Blast | 2.9 |
| tRNA prediction | tRNAscan-SE | 2.05 |
| **Mitogenome assembly** |  |  |
| Read alignment | Minimap2 | 2.24 |
| Read extraction | Samtools | 1.16.1 |
| Assembly | Canu | 2.1.1. |
| Visual inspection and consensus | Geneious | 11.0.3 |
| Annotation | MitoZ | 3.4 |
| **Genome quality assessment** |  |  |
| General metrics | QUAST | 5.2.0 |
| Genome quality and completeness | BUSCO | 4.0.5 |

## Results

### Nuclear assembly

We generated a de novo nuclear genome assembly of the northern bat (*BLF_Eptnil_asm_v1*.*0*) using 12 million PacBio CLR reads and 160 million read pairs of OmniC data. The Pacbio data yielded ∼141 fold coverage (N50 read length 31,406 bp; minimum read length 50 bp; mean read length 22,041.9 bp; maximum read length 27,6123 bp) based on the final assembled genome size of 2.0 Gb. Assembly statistics are reported in tabular form in Table 2. The final assembly consists of 726 scaffolds spanning 2.0 Gb with contig N50 of 66.2 Mb, scaffold N50 of 142.1 Mb, largest contig of 202.7 Mb, largest scaffold of 239.8 Mb and number of gaps is 166. The Omni-C contact map suggests that the primary assembly is highly contiguous (Figure 2). Gene annotation predicted total of 20 250 genes. RepeatMasker masked 36.41% of the genome of which class I TEs repeats were 18.17% and class II TEs repeats 2.98%. The assembly has a BUSCO completeness score of 93.73% using the eukaryota gene set.

**Table 2.**
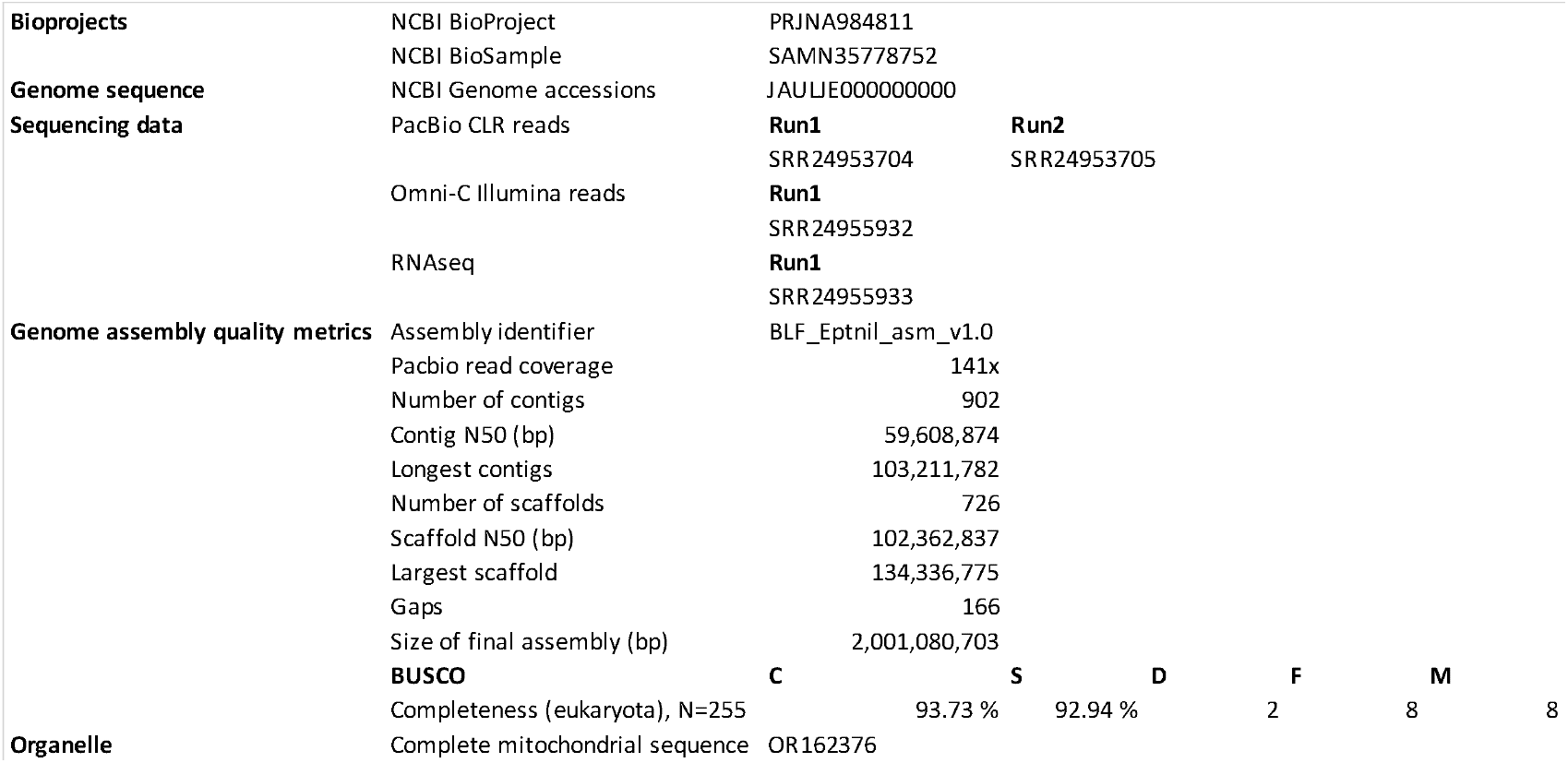
Summary statistics of the sequencing datasets used and the assembly.

**Figure 2.**
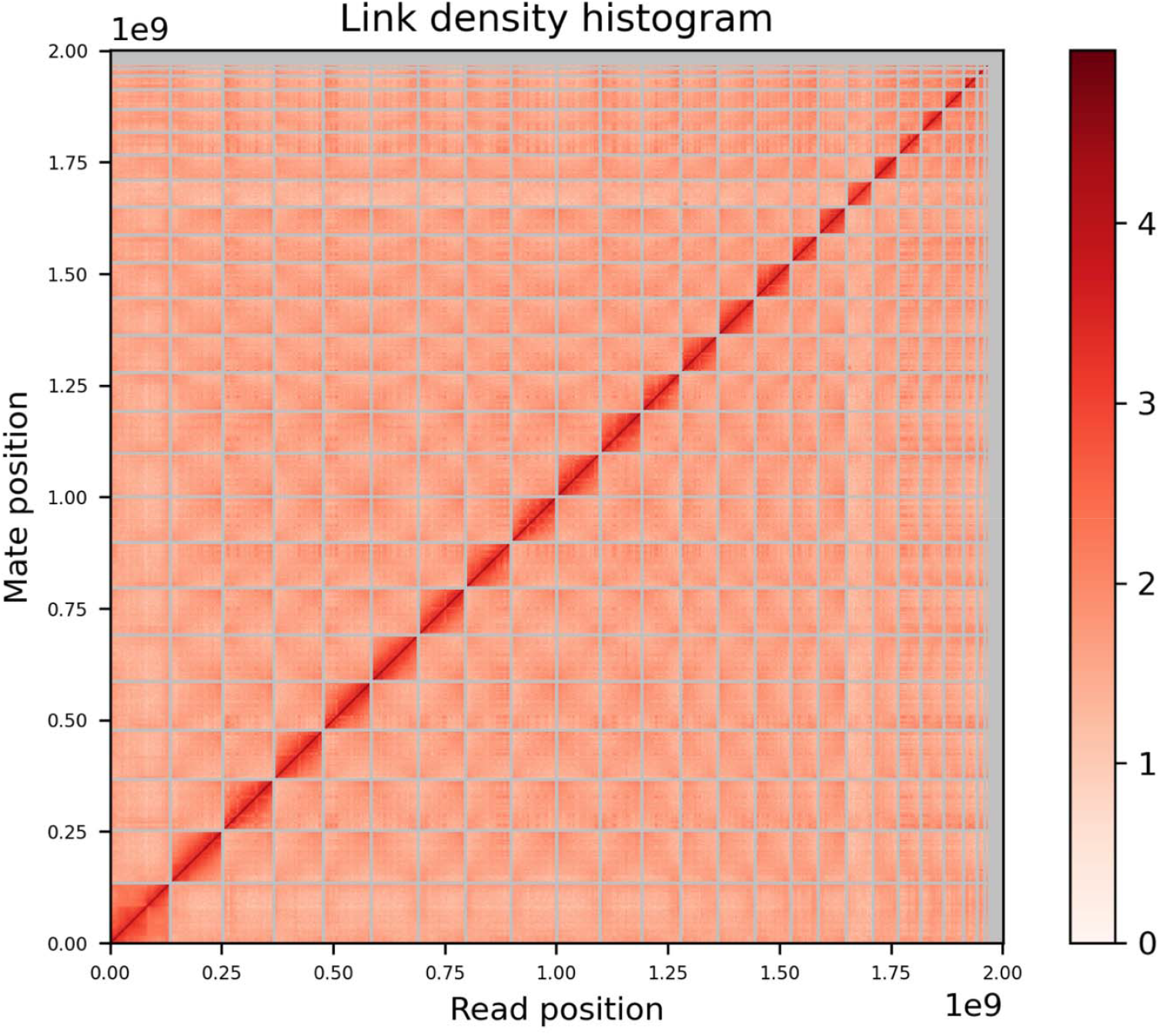
Omni-C contact density map.

### Mitochondrial assembly

The mitochondrial genome assembled with Canu resulted in five contigs which were aligned with Geneious and the final consensus had size of 17 011 bp. The base composition of the final assembly version is A = 32.1%, C = 23.9%, G = 14.9%, T = 29.1%, and consists of 22 transfer RNAs and 13 protein coding genes.

## Discussion

Here, we present high-quality reference genome for *Eptesicus nilssonii*. It adds to the growing number of bat genomes available for research (Jebb et al. 2020). The production of this reference genome can assist in understanding the fate of the species by allowing in-depth analysis of population connectivity and structure, the number of ancestral populations and possible population differentiation that could have led to potentially beneficial local adaptations. Furthermore, the production of various genomes assists in illuminating the spectacular adaptive radiation and systematics of bats (Jebb et al., 2020), with further potential of uncovering mechanisms that allow bats to harbor a variety of zoonotic pathogens without apparent harm (Jakava-Viljanen et al., 2010; Kivistö et al., 2019; O’Shea et al., 2014; Veikkolainen et al., 2014).

*E. nilssonii* has a remarkable distribution range spanning the entire Palearctic boreal zone (Suominen et al., 2020). However, the species also has little possibility adjust its distribution range to the north in response to the current climate change. With *E. nilssonii* being the northernmost bat species in the world, we can also expect the reference genome to provide additional and complimentary insights into genetic components of cold adaptation (Yurchenko et al. 2018).

We also broaden the potential use of cell lines to the assist in the production of high-quality reference genomes. Genomes, such as the one presented here provide researchers and conservation scientists tools to access the most advanced downstream bioinformatic methods to better safeguard global biodiversity.

## Acknowledgements

No ethics permits were needed in the process. We thank Dovetail Genomics for the sequencing, help and support.

## Data Availability

Data generated for this study are available under NCBI BioProject PRJNA984811. Raw sequencing data for reference sample (NCBI BioSample SAMN35778752) are deposited in the NCBI Short Read Archive (SRA) under SRR24953704 and SRR24953705 for PacBio sequencing data, SRR24955932 for Omni-C Illumina Short read sequencing data and for RNAseq SRR24955933. GenBank accessions for the assembly is JAULJE000000000. The GenBank organelle genome assembly for the mitochondrial genome is OR162376.

## Supplementary Material

**Supplementary Figure 1**. Morphology of the clonal SV40LT-immortalized kidney cells of the reference northern bat. Phase contrast microscope images at 40x of all the isolated clonal cell lines. The clonal isolate 8+ was used for the sequencing.

